# The Nucleocapsid Protein of SARS-CoV-2 Abolished Pluripotency in Human Induced Pluripotent Stem Cells

**DOI:** 10.1101/2020.03.26.010694

**Authors:** Zebin Lin, Zhiming Wu, Jinlian Mai, Lishi Zhou, Yu Qian, Tian Cai, Zhenhua Chen, Ping Wang, Bin Lin

**Author notes:** These authors contributed equally to the work (Zebin Lin, Zhiming Wu). **Correspondence:** Ping Wang; Bin Lin.

## Abstract

The COVID-19 pandemic caused by severe acute respiratory syndrome coronavirus 2 (SARS-CoV-2) is raging across the world, leading to a global mortality rate of 3.4% (estimated by World Health Organization in March 2020). As a potential vaccine and therapeutic target, the nucleocapsid protein of SARS-CoV-2 (nCoVN) functions in packaging the viral genome and viral self-assembly. To investigate the biological effects of nCoVN to human stem cells, genetically engineered human induced pluripotent stem cells (iPSC) expressing nCoVN (iPSC-nCoVN) were generated by lentiviral expression systems, in which the expression of nCoVN could be induced by the doxycycline. The proliferation rate of iPSC-nCoVN was decreased. Unexpectedly, the morphology of iPSC started to change after nCoVN expression for 7 days. The pluripotency marker TRA-1-81 were not detectable in iPSC-nCoVN after a four-day induction. Meanwhile, iPSC-nCoVN lost the ability for differentiation into cardiomyocytes with a routine differentiation protocol. The RNA-seq data of iPSC-nCoVN (induction for 30 days) and immunofluorescence assays illustrated that iPSC-nCoVN were turning to fibroblast-like cells. Our data suggested that nCoVN disrupted the pluripotent properties of iPSC and turned them into other types of cells, which provided a new insight to the pathogenic mechanism of SARS-CoV-2.

## 1 Introduction

Right now, the COVID-19 pandemic is sweeping the world, causing a huge crisis in public health and economics globally. According to the continuously updated data from World Health Organization, to date, nearly three million infected cases were confirmed, while more than 200,000 individuals died because of COVID-19 (https://www.who.int/emergencies/diseases/novel-coronavirus-2019). Severe acute respiratory syndrome coronavirus 2 (SARS-CoV-2), which was proved to be the pathogen of COVID-19, has 79% identity in genomes with severe acute respiratory syndrome coronavirus (SARS-CoV) (Lu et al., 2020). Twelve coding regions were predicted in SARS-CoV-2, including spike protein, nucleocapsid protein, envelope protein, and membrane protein (Lu et al., 2020; Wu et al., 2020; Xu et al., 2020). The Cryo-EM structure of spike protein had been determined (Wrapp et al., 2020), and more and more evidences showed that the spike protein binds human ACE2 to entry into host cells (Hoffmann et al., 2020; Wrapp et al., 2020), which indicated that SARS-CoV-2 might share similar pathogenic mechanisms with SARS-CoV. Because of the very limited knowledge of SARS-CoV-2, we sought to understand the biology of SARS-CoV-2 based on the previous studies about SARS-CoV.

As one of the most studied proteins in SARS-CoV, the nucleocapsid protein binds to viral RNA to package the genome in a ribonucleoprotein particle (Chang et al., 2014). Unlike the spike protein with a certain mutation frequency, the sequence of nucleocapsid protein was more stable (Chinese, 2004), which meant it was an ideal target for diagnostic tools (Severance et al., 2008; Suresh et al., 2008; Das et al., 2010) and antiviral therapy (Cheung et al., 2008; Chang et al., 2016). The pathogenic effects in host cells caused by the nucleocapsid protein were also studied. It was reported that the nucleocapsid protein inhibited type I interferon production after virion infected the host cells (Hu et al., 2017), which was considered as a possible mechanism of immune escape. The nucleocapsid protein inhibited cell cytokinesis and proliferation (Zhou et al., 2008), and regulated several pathways, such as transforming growth factor-beta signaling (Zhao et al., 2008), AP-1 signal transduction pathway (He et al., 2003), and NF-KappaB pathway (Zhang et al., 2007b). Besides, the nucleocapsid protein was reported as an apoptosis inducer in COS-1 cells (Surjit et al., 2004; Zhang et al., 2007a) and HPF cells (Zhao et al., 2006).

As the nucleocapsid protein of SARS-CoV-2 (nCoVN) has 88.1% identity with the nucleocapsid protein of SARS-CoV (Lu et al., 2020), it is reasonable to speculate that they share a same pathogenic pathway in host cells. The original goal of this study is to determine the physiological malfunctions, such as cardiac fibrosis, in human cardiomyocytes expressing nCoVN by using human induced pluripotent stem cells (iPSC) and direct cardiac-differentiation methods. However, the morphology of iPSC altered obviously when nCoVN had been expressed for 7 days. This unexpected observation inspired us some new thoughts: (i) It was likely that the adult stem cells could be infected by SARS-CoV-2, in spite of lacking of the clinical data; (ii) iPSC were appropriate study materials for stem cell research because of avoiding many ethical issues; (iii) The preliminary data indicated that nCoVN seemed to be deleterious to iPSC, which meant it might also cause damages in other stem cells. Therefore, we turned to investigate whether nCoVN obstructed the pluripotency maintenance in iPSC. We believed that this study could help us to understand the deleterious effects of nCoVN to human adult stem cells and embryonic stem cells.

## 2 Method

### 2.1 Cell culture and differentiation assay

Human induced pluripotent stem cells (iPSC) DYR0100 (from The American Type Culture Collection, ATCC) were plated on Matrigel matrix (hESC-Qualified, LDEV-Free, Corning, 354277)-coated plates, and then were cultured in DMEM/F-12 medium (Gibco, 11320033) supplemented with STEMUP^®^ ES/iPS cell culture medium supplement (Nissan Chemical Corporation). STEMUP medium was changed every two days. iPSC were passaged every three to four days or when the cell culture was 80-90% confluent. During passages, iPSC were rinsed with 1× DPBS (Gibco, 14040133) for one time then were treated with 0.5 mM EDTA (Invitrogen, 15575020) in 1× DPBS (Gibco, 14190144) for 10 minutes at room temperature. The split ratio was 1:3-1:6. The detailed differentiation protocol was described in the previous published reports (Lin et al., 2017; Shekhar et al., 2018). Briefly, iPSC were treated with the small molecule CHIR99021 (Tocris, 4423, final concentration 10 μM) in the RPMI-BSA medium [RPMI 1640 Medium (HyClone, SH30027.01) supplemented with 213 μg/ml AA2P (l-ascorbic acid 2-phosphate magnesium) (Sigma, A8960) and 0.1% bovine serum albumin (BSA) (Sigma, A1470)] for 24 hours, then were incubated with RPMI-BSA medium for 48 hours. On differentiation day 4, cells were treated with the small molecule IWP2 (Tocris, 3533, final concentration 5 μM) in RPMI-BSA medium. After 48 hours, the medium was changed to RPMI-BSA medium. Then, RPMI 1640 medium supplemented with 3% KnockOut Serum Replacement (Gibco, 10828-028) was used to culture the cardiomyocytes in the following experiments. All the cells in this study (except iPSC-derived cardiomyocytes) were kept culturing in the STEMUP medium until they were applied to other assays.

### 2.2 Generation of iPSC-nCoVN

The cDNA of nCoVN with a N-terminal 6× His Tag coding sequence (GeneMedi) and puromycin resistance gene were sub-cloned into the plasmid pCW-Cas9-Blast (Addgene, 83481) to replace Cas9 and Blast cDNA, respectively. Lentivirus preparation using a third generation lentivirus packaging system were referred to the previous report (Jiang et al., 2015). We followed and modified the protocol from Zhang lab to detect MOI of the lentivirus and perform transduction (Shalem et al., 2014). After 24 hours of transduction, medium was changed to fresh STEMUP medium supplemented with doxycycline hyclate (Sigma, D9891) for induction. Two days later, puromycin (InvivoGen, ant-pr-1, final concentration 2 μg/mL) was added into the STEMUP medium supplemented with doxycycline hyclate. After 2-3 days' selection, which resulted in a transduction efficiency of ~30%, single cell clones were manually picked and re-seeded in separated wells. For nCoVN expression induction, doxycycline hyclate (Sigma, D9891) was supplemented in the stem cell culture medium at a final concentration of 2 μg/mL, and the same amount of DMSO was added to the stem cell culture medium for controls.

### 2.3 Reverse transcription-PCR and Quantitative Real-time PCR

Total RNA was extracted by using the UNlQ-10 Column Trizol Total RNA Isolation Kit (Sangon Biotech, B511321-0100) prior to the treatment with DNase I (Sangon Biotech, B618252) for 30 minutes. mRNA was reverse transcribed by using iScript Reverse Transcription Supermix (Bio-Rad, 1708841). Quantitative Real-time PCR was performed by using a PikoReal Real-Time PCR System (Thermo Fisher) with SsoAdvanced™ Universal SYBR^®^ Green Supermix (Bio-Rad, 1725271). The primers for Reverse transcription-PCR and Quantitative Real-time PCR are as followed (from 5′ to 3′):

ACE2-RT-F: GGTCTTCTGTCACCCGATTT;
ACE2-RT-R: ACCACCCCAACTATCTCTCG;
nCoVN-RT-F: CATTGGCATGGAAGTCACAC;
nCoVN-RT-R: TCTGCGGTAAGGCTTGAGTT;
GAPDH-RT-F: TGGGTGTGAACCATGAGAAG;
GAPDH-RT-R: GTGTCGCTGTTGAAGTCAGA.

### 2.4 The proliferation assay

iPSC, iPSC-GFP and iPSC-nCoVN were seeded in 96-well plates with the same cell number. After 24 hours, CCK-8 reagent was added in the medium to monitor the proliferation rate (Beyotime, C0038). The absorbance at 450 nm was measured at 24 hours, 42 hours, 48 hours, 60 hours and 72 hours by using Varioskan Flash Multimode Reader (Thermo Scientific). The cell-free medium with CCK-8 reagent were used as blank control sets. The data were analyzed and plotted using GraphPad Prism 6.

### 2.5 Immunofluorescence Staining

Cells were fixed with 4% paraformaldehyde at room temperature for 20 minutes and washed three times with 1× PBS. Cells were then permeabilized with PBS containing 0.25% Triton X-100 at room temperature for 10 minutes. After incubating in the blocking buffer (1× PBS with 10% goat serum), cells were stained with different primary antibodies at 4°C overnight. These primary antibodies were [target, dilution, species, company, product number]: Troponin T Cardiac Isoform, 1:100, mouse, Thermo Fisher, MA5-12960; alpha-smooth muscle actin, 1:100, mouse, Bioss, bsm-33187M; S100A4, 1:100, rabbit, Bioss, bs-3759R; SSEA4, 1:250, mouse, Invitrogen, 14-8843-80; TRA-1-81, 1:250, mouse, Invitrogen, 14-8883-80; 6× His Tag, 1:250, mouse, Sangon Biotech, D191001; OCT4, 1:200, mouse, Abcam, ab184665; ACE2, 1:200, rabbit, Bioss, bs-1004R; vimentin, 1:250, rabbit, Bioss, bs-0756R. Cells were washed three times with PBS containing 0.1% Triton X-100, then incubated with the Alexa Fluor 488 goat anti-mouse or Alexa Fluor 555 goat anti-rabbit IgG secondary antibodies at 37°C for 1 hour. Nuclei were labeled with DAPI (4′,6-diamidino-2-phenylindole, 1 μg/ml) for 5 minutes. Images were obtained by using the DMi6000 B inverted microscope (Leica) or the FV1000 confocal laser scanning microscope (Olympus), then were analyzed by using ImageJ software.

### 2.6 RNA-seq analysis method

RNA-seq was performed by Novogene Co., Ltd. We obtained 151 bp paired-end RNA-seq reads from an Illumina Novaseq instrument, average 23 million read pairs for 3 iPSC-GFP and 3 iPSC-nCoVN samples. Adapters and low-quality bases in reads were trimmed by trim_galore (v0.6.5; http://www.bioinformatics.-babraham.ac.uk/projects/trim_galore/). We employed Kallisto (v0.46.0) (Bray et al., 2016) to determine the read count for each transcript and quantified transcript abundance as transcripts per kilobase per million reads mapped (TPM), using gene annotation in the GENCODE database (v32, GRCh38) (Frankish et al., 2019). Then we summed the read counts and TPM of all alternative splicing transcripts of a gene to obtain gene expression levels. We restricted our analysis to 22,201 expressed genes with an average TPM >=1 in either iPSC-GFP or iPSC-nCoVN samples. DESeq2 (v1.26.0) (Love et al., 2014) was used to identify differentially expressed genes (DEGs) (false discovery rate (FDR) <0.05 and abs(log2FoldChange) >3). The pathway analysis was performed by ToppGene Suite (Chen et al., 2009). The raw RNA-seq data have been deposited in GEO, and the accession number is pending.

We took gene expression values (i.e. log_2_(TPM)) in iPSC/ESC and fibroblast from ENCODE (Consortium, 2012; Davis et al., 2018). Combining with our RNA-seq data, quantile normalization (Bolstad et al., 2003) was performed, and then ComBat (Johnson et al., 2007) was used to remove the batch correction. We selected the top 1000 genes with the largest variance to calculate the correlation coefficients between samples. The heatmap was generated by pheatmap function in R.

### 2.7 Statistic

Values were expressed as mean ± SD (standard deviation). Statistical significances were evaluated using one-way ANOVA with Bonferroni correction or Student’s T-Test. *P*<0.05 was considered statistically significant.

## 3 Results

### 3.1 *ACE2* was expressed in various of stem cells

As ACE2 was the major receptor of SARS-CoV-2 on the cell membrane (Hoffmann et al., 2020; Wrapp et al., 2020), we first examined whether *ACE2* was expressed in the stem cells. Thanks to the gene expression data collection in Gene Expression Omnibus (GEO, https://www.ncbi.nlm.nih.gov/geo/), it was convenient to analyze the *ACE2* expression profiles in sorts of stem cells. Figure 1A showed the *ACE2* expression values in different stem cells from different projects, including human embryonic stem cells (Kim et al., 2014), iPSC (Yang et al., 2014), human epithelial stem cells (Yang et al., 2014), human adipose stem cells (Onate et al., 2013), human hematopoietic stem cells (Pang et al., 2011), and human mesenchymal stem cells (Bernstein et al., 2010). The expression values of a housekeeping gene *GAPDH* were simultaneously collected as controls. *ACE2* was expressed in each kind of stem cells, though the expression values were relatively low compared with *GAPDH*. The reverse transcription-PCR (RT-PCR) results showed that *ACE2* was expressed in iPSC, iPSC-derived cardiomyocytes (iPSC-CM) and human coronary artery endothelial cells (HCAEC) (Figure 1B). The images from immunofluorescence assays clearly showed that ACE2 protein was located on the cell membrane of iPSC (Figure 1C), suggesting that the pluripotent stem cells were the potential targets of SARS-CoV-2.

**Figure 1.**
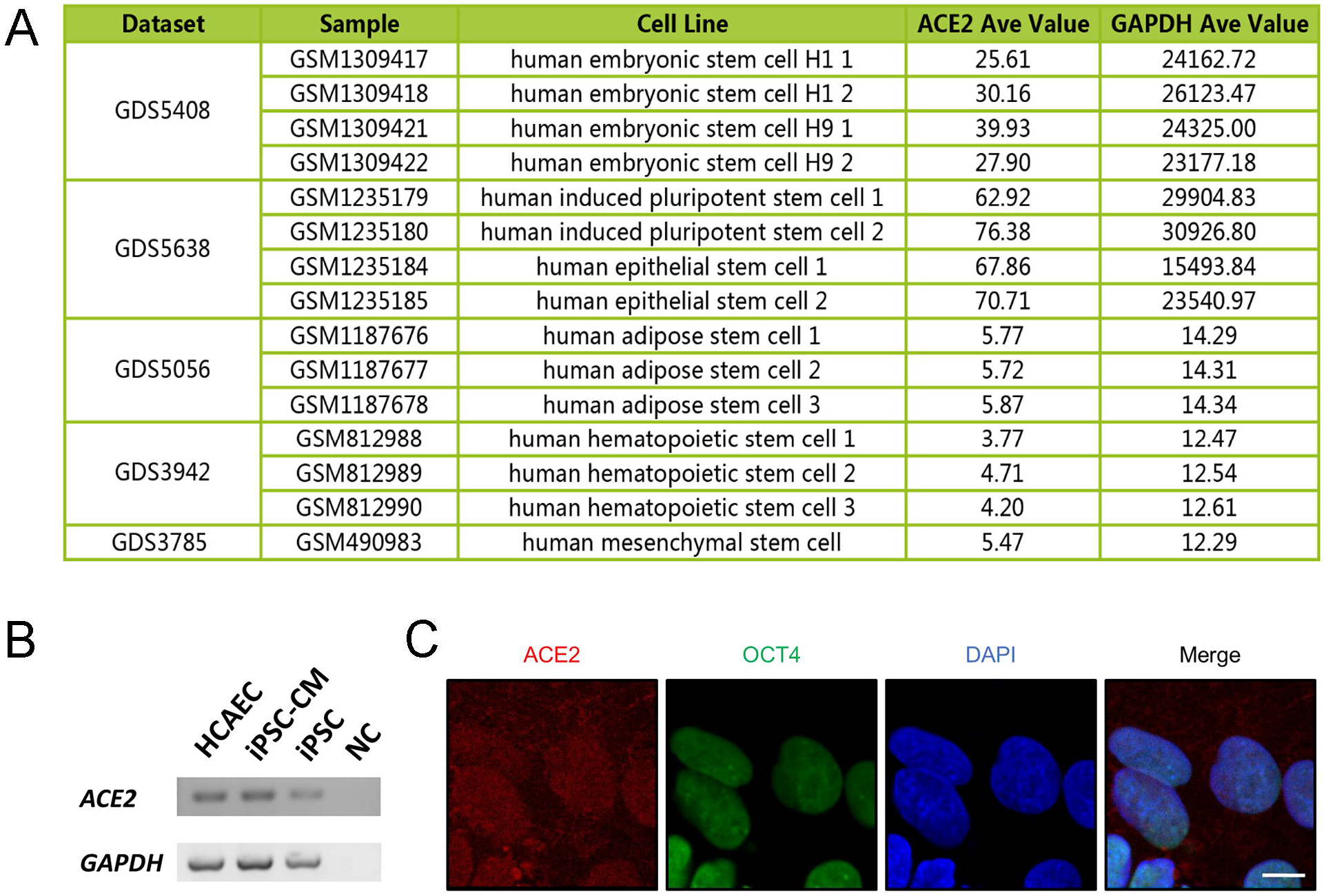
*ACE2* was expressed in human stem cells. (A) Expression values of *ACE2* and *GAPDH* derived from the Gene Expression Omnibus database. (B) Images from agarose gel electrophoresis for analyzing the Reverse transcription-PCR products. *ACE2* was expressed in iPSC, iPSC-CM and HCAEC. iPSC, human induced pluripotent stem cell; iPSC-CM, human induced pluripotent stem cell-derived cardiomyocyte; HCAEC, human coronary artery endothelial cell; NC, negative control. (C) Representative immunofluorescent staining images of ACE2 (red) and the pluripotency marker OCT4 (green) in iPSC. The cell nuclei were stained by DAPI (blue). The scale bar represents 10 μm.

### 3.2 Expression of nCoVN changed the morphology of iPSC

To study whether physiological activities in iPSC were disturbed by nCoVN, a human induced pluripotent stem cell line (iPSC-nCoVN) in which the expression of nCoVN could be modulated by a Tet-On system was generated by a lentiviral expression system. In this system, nCoVN cDNA sequence (with a 6× His Tag coding sequence) was conjugated to puromycin resistance gene through a T2A peptide encoding sequence, and the transcription was relied on the induction of tetracycline or doxycycline (Dox). After puromycin selection, two single cell clones were seeded in separated wells by manual colony-picking. Sequentially, iPSC-nCoVN were divided into two groups: one was induced by Dox for nCoVN expression (Dox), the other was added with DMSO as a control set (DMSO), meanwhile, a GFP-expressed iPS cell line (iPSC-GFP), in which the expression of GFP was modulated by the same Tet-On system, was used as another control set in the following assays (Figure 2A).

**Figure 2.**
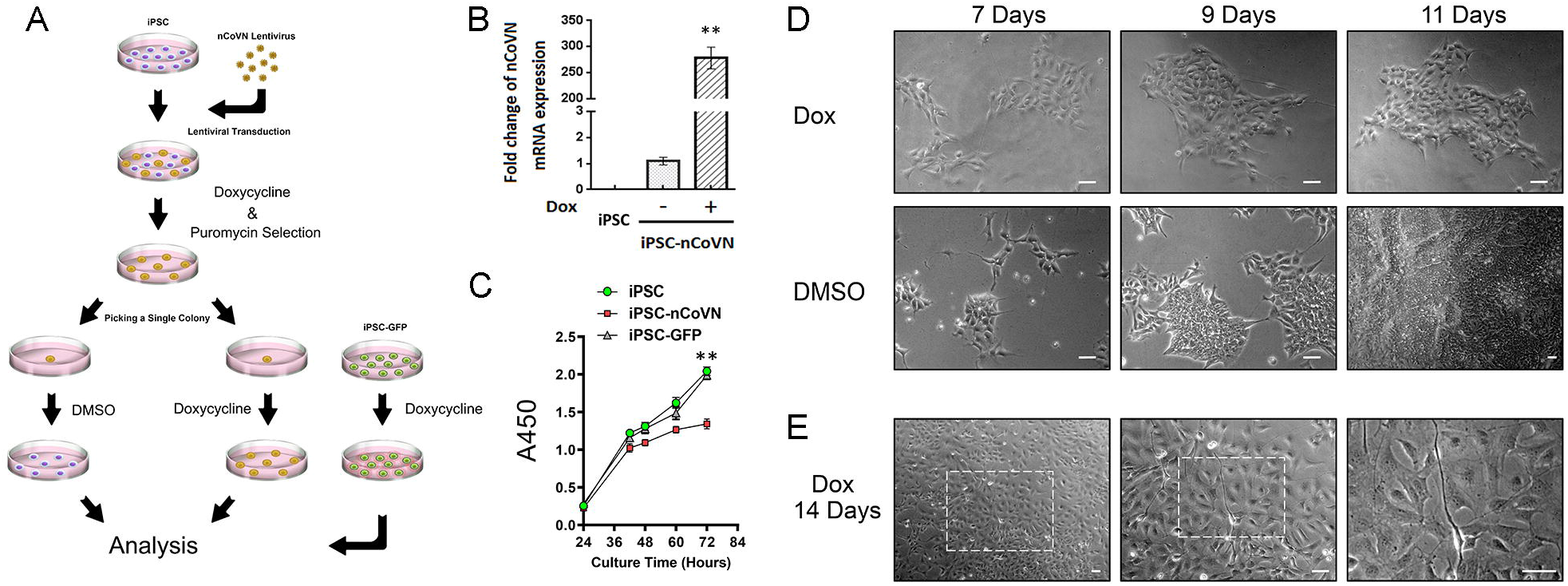
nCoVN affected the proliferation and morphology of iPSC. (A) Schematic diagram illustrating the generation of iPSC-nCoVN and controls. Purple cells indicated iPSC without nCoVN expression, while yellow cells indicated iPSC with nCoVN expression. Green cells were iPSC stably expressing GFP under the doxycycline induction. (B) The mRNA expression level of nCoVN was significantly elevated in iPSC-nCoVN for a long-term induction (n=3). **, *p*<0.001. (C) The time course of cellular proliferation from iPSC, iPSC-GFP and iPSC-nCoVN (n=6). At 72 hours after cell seeding, the values of A450 were significantly increased in iPSC and iPSC-GFP groups. **, *p*<0.001. (D) Representative phase-contrast images from nCoVN-positive cells (Dox group) and control cells (DMSO group) after a 7-day, a 9-day, and an 11-day inductions. The scale bar is 50 μm. (E) Representative phase-contrast images of iPSC-nCoVN after a 14-day induction show detailed morphological alterations. Images were taken under objectives with 10×, 20×, and 40× magnifications (from left to right panels). White dashed line boxes indicate the regions that are magnified in the right panel. The scale bar represents 50 μm.

The expression of nCoVN was confirmed at the mRNA and protein levels. The transcriptional level of nCoVN was measured by Real-time PCR in iPSC, Dox, and DMSO groups, and nCoVN expression increased about 267-fold in the Dox group compared with the DMSO group (Figure 2B). The nCoVN protein was detected by using an anti-6× His Tag antibody in cells from the Dox group (Supplementary Figure 1). The proliferation rate was compared among iPSC, iPSC-GFP and Dox groups by using a cell counting kit. The absorbance at 450 nm (A450) was measured at 24 hours, 42 hours, 48 hours, 60 hours and 72 hours after cell seeding. After three days of cell seeding, Dox group showed a decreased proliferation rate than both of iPSC and iPSC-GFP groups, indicating that nCoVN might hamper the growth and division of iPSC (Figure 2C). This observation was consistent with the previous finding about the nucleocapsid protein of SARS-CoV (Zhou et al., 2008).

We continued to induce nCoVN expression in iPSC. The phase-contrast images of iPSC-nCoVN with a 7-day, a 9-day, and an 11-day inductions (Dox) and counterpart controls (DMSO) were shown in Figure 2D. In the DMSO group, a typical morphology of stem cells with high nucleus/cytoplasm ratio and close cell membrane contacts was observed, while iPSC-nCoVN after a 7-day induction started to exhibit endothelial cell morphological features and lower nucleus/cytoplasm ratio. After a 14-day induction, most of the cells exhibited distinct shapes from wild-type iPSC, such as neuron-like cells, endothelial-like cells and fibroblast-like cells (Figure 2E). These data showed that continuous expression of nCoVN caused obvious morphological changes in iPSC.

### 3.3 Expression of nCoVN disabled the pluripotent properties of iPSC

Next, we examined the pluripotency markers in iPSC and iPSC-nCoVN. The pluripotency markers SSEA4 and TRA-1-81, which were expressed in human embryonic stem cells and iPSC, were widely applied in identification of pluripotent stem cells (Abujarour et al., 2013; Trusler et al., 2018). The immunofluorescence staining images illustrated that iPSC-nCoVN completely lost the expression of SSEA4 and TRA-1-81, namely, iPSC-nCoVN lost the pluripotency in the presence of nCoVN (Supplementary Figure 2A, B). We traced the expression of TRA-1-81 in iPSC-nCoVN with a 2-day, a 4-day, a 6-day, and an 8-day inductions (Figure 3A). On Day 2, TRA-1-81 was still expressed in iPSC-nCoVN; however, from Day 4, TRA-1-81 was not detectable in most of the cells, suggesting that the pluripotent fate of iPSC-nCoVN was determined in the first 4 days. To further test the pluripotency in iPSC and iPSC-nCoVN, we directly differentiated these cells to cardiomyocytes by using a routine protocol, and the differentiation assays were performed under the same conditions. As expected, the differentiation efficiency could reach 60% in iPSC; however, on differentiation day 12, only a very small portion of cells from iPSC-nCoVN were expressed cardiac Troponin T, accompanied by many cell deaths (Figure 3B, C). This differentiation assay provided solid evidence that the pluripotency maintenance of iPSC-nCoVN was disrupted by nCoVN.

**Figure 3.**
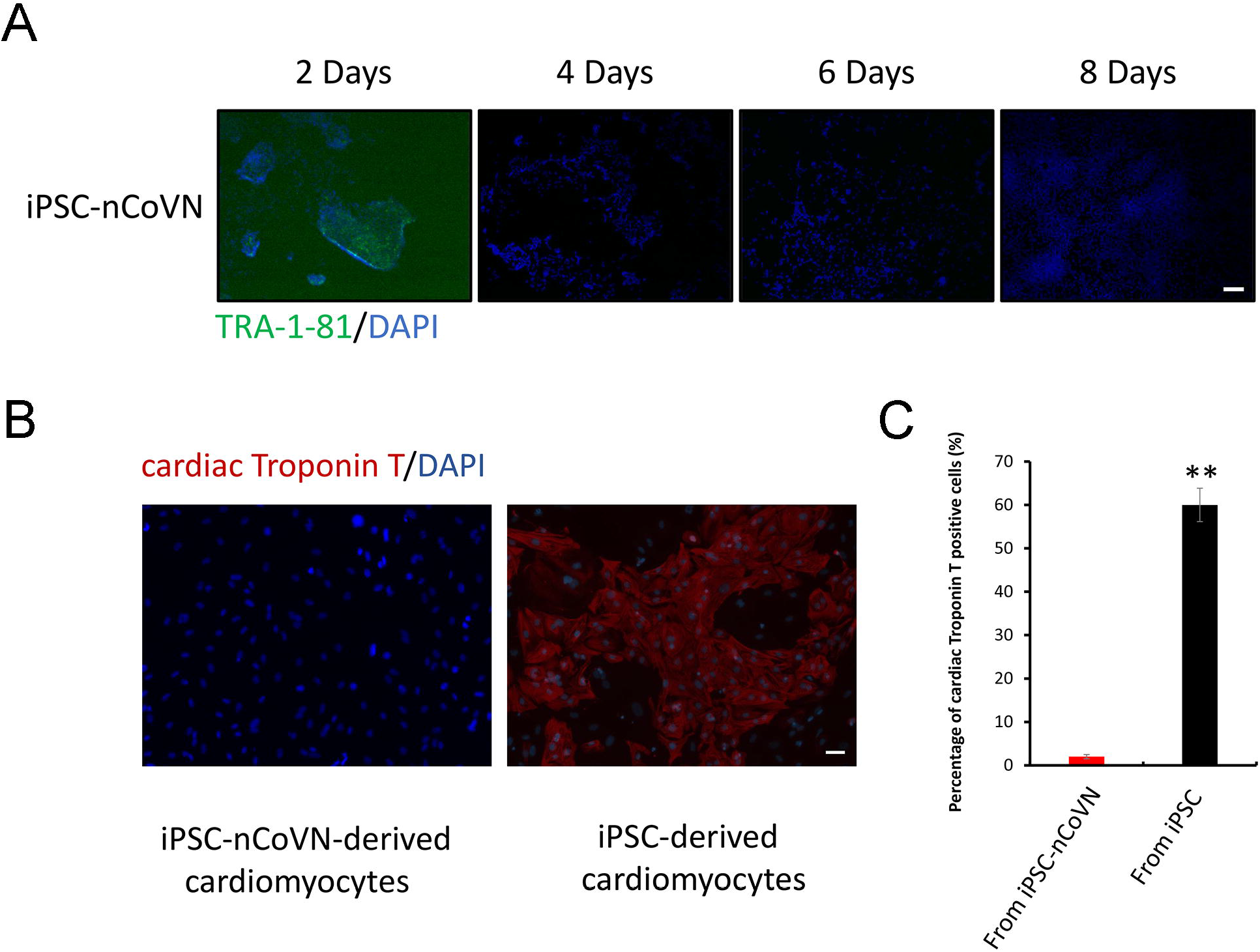
iPSC-nCoVN lost the pluripotency. (A) Representative immunofluorescent staining images of pluripotency marker TRA-1-81 (green) in iPSC-nCoVN for a 2-day, a 4-day, a 6-day, and an 8-day inductions. The cell nuclei were stained by DAPI (blue). The scale bar represents 50 μm. (B) Representative immunofluorescent staining images of cardiomyocyte marker cardiac Troponin T (Red) in iPSC- and iPSC-nCoVN-derived cardiomyocytes. The cell nuclei were stained by DAPI (blue). The scale bar represents 50 μm. (C) The cardiac differentiation efficiency of iPSC and iPSC-nCoVN. Images taken from (B) were analyzed by using ImageJ software. The efficiency was calculated as the portion of cardiac Troponin T positive cells in all the cells. Approximately 6,000 cells were counted in each group. **, *p*<0.001.

### 3.4 Long-term expression of nCoVN drove iPSC to fibroblast

Since the pluripotency lost due to short-term expression of nCoVN, we are extremely interested in the cell fate of iPSC-nCoVN under long-term expression of nCoVN. After a 28-day induction in the stem cell culture medium, some spindle-shaped iPSC-nCoVN, which exhibited a typical fibroblast morphological feature, were observed (Supplementary Figure 2C). The antibodies against fibroblast markers vimentin, alpha-smooth muscle actin (α-SMA) and S100A4 were used to verify the cell type of these fibroblast-like cells. The results from immunofluorescence assays confirmed that these markers were expressed in nCoVN-expressing cells (Figure 4A, B, C; Supplementary Figure 2D). To further investigate the transcriptomic profiles of iPSC-nCoVN under the long-term nCoVN expression, doxycycline-induced iPSC-nCoVN and iPSC-GFP for 30 days were applied to RNA-seq. Through differentially express analysis, iPSC-nCoVN showed a dramatic gene expression change comparing with iPSC-GFP (Supplementary Table). Totally, 3,080 genes were significantly differentially expressed (FDR<0.05, |log2FoldChange|>3). Among them, the down-regulated genes in iPSC-nCoVN were most significantly enriched with proliferation and stem cell related pathways, including the Yamanaka factors-associated genes, such as *POU5F1*, *LIN28A*, *NANOG*, and *SOX2* (with a 790-fold, a 2306-fold, a 253-fold, and an 18-fold decrease, respectively) (Figure 4E); while the extracellular matrix and extracellular matrix-associated pathway was the most significantly enriched pathway in the up-regulated genes (Figure 4F). Next, we used RNA-seq data from the ENCODE project to evaluate the cell type of iPSC-nCoVN. Comparing with the transcriptome of the pluripotent stem cells and fibroblast in the ENCODE project (Consortium, 2012; Davis et al., 2018), iPSC-GFP samples were clustered with H7 and GM23338, which were the embryonic stem cells (ESC) and iPSC, respectively; while iPSC-nCoVN samples were clustered with multiple kinds of fibroblast (Figure 4G). Furthermore, iPSC-nCoVN with a 40-day induction, which were kept culturing in the stem cell medium, were totally differentiated to fibroblast (Figure 4D).

**Figure 4.**
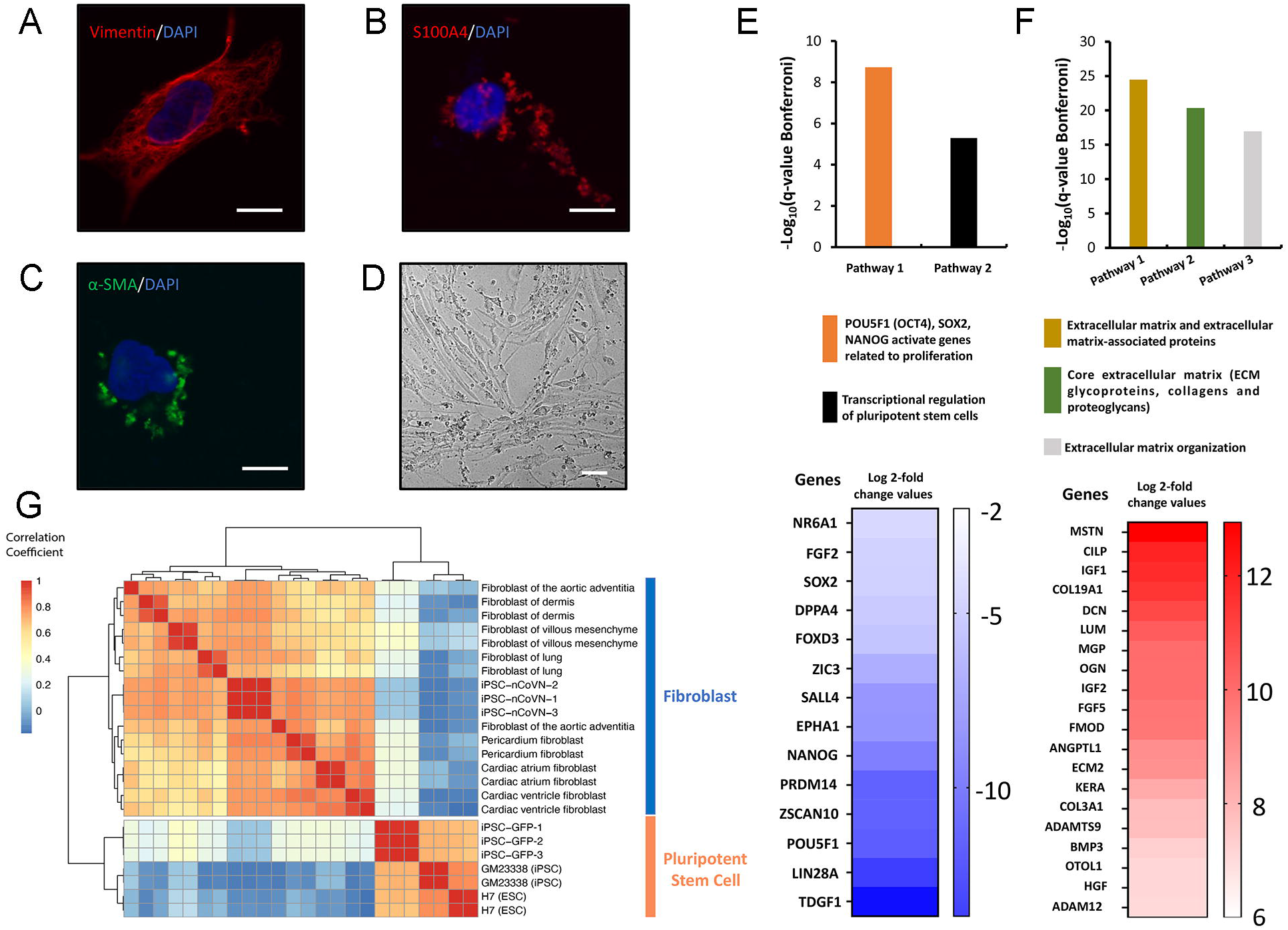
Long-term expression of nCoVN turned iPSC to fibroblast. (A-C) Representative immunofluorescent staining images of vimentin (green), S100A4 (green), and α-SMA (green) in iPSC-nCoVN after a 10-day induction. The cell nuclei were stained by DAPI (blue). The scale bars represent 10 μm. (D) The representative bright field image of cells after a 40-day nCoVN expression. The morphology exhibits typical fibroblast features. The scale bar represents 50 μm. (E) The two most significantly enriched pathways in the down-regulated genes. The histogram shows the significance of the two pathways by using −log_10_(q values with Bonferroni correction). The Log 2-fold change values of a total of 14 genes in these two pathways were exhibited by the heatmap. (F) The three most significantly enriched pathways in the up-regulated genes. The histogram shows the significance of the three pathways by using −log_10_(q values with Bonferroni correction). The Log 2-fold change values of the top 20 up-regulated genes in these pathways were exhibited by the heatmap. (G) The heatmap of the correlation coefficients among iPSC-GFP, iPSC-nCoVN, ESC/iPSC and fibroblast from the ENCODE project. The 1000 most variable genes in the samples were used to calculate the correlation coefficients. iPSC-GFP and iPSC-nCoVN clustered with pluripotent stem cell and fibroblast, respectively.

## 4 Discussion

According to the current knowledge about the life cycle of SARS-CoV, the nucleocapsid protein was translated by the host cell translation protein synthesis machinery (McBride et al., 2014; Song et al., 2019), and was localized mainly in the cytoplasm (Rowland et al., 2005). The primary function of nucleocapsid protein was to package the viral genome into nucleocapsids to protect the genomic RNA (McBride et al., 2014). During the formation of nucleocapsids, numerous nucleocapsid proteins bound to the viral RNA and started oligomerization. The viral reproductive strategies would synthesize nucleocapsid proteins as many as possible to meet the requirements of viral assembly, which meant the nucleocapsid proteins were overproduced. The findings that redundant nucleocapsid proteins interfered with the normal physiology of host cells were reported (Surjit et al., 2004; Zhao et al., 2006; Zhang et al., 2007a; Zhao et al., 2008; Zhou et al., 2008; Hu et al., 2017). In this study, we first presented that nCoVN abolished pluripotency and reduced the proliferation rate in human induced pluripotent stem cells. Long-term expression of nCoVN drove iPSC to fibroblast in spite of using the stem cell culture conditions. It was reported that the nucleocapsid protein of SARS-CoV facilitated TGF-β-induced PAI-1 expression to promote lung fibrosis (Zhao et al., 2008), which was also the possible pathway that nCoVN turned iPSC to fibroblast.

The time-course assays showed that the pluripotency marker disappeared in four days after nCoVN expression. This finding might be applied to a cell-based chemical screening model, in which the candidate chemicals with potential ability to halt the iPSC differentiation caused by nCoVN are easily identified. More importantly, SARS-CoV-2 is not necessary in this model, which means it could be used in the routine laboratories and applied to high-throughput equipment with less risk.

In addition, how nCoVN breaks the pluripotency maintenance of iPSC is still a riddle. The pluripotency maintenance in stem cells requires delicate regulations to maintain the balance of pluripotency gene expression in a complicated network. Since nCoVN can bind RNAs, it is possible that nCoVN suppresses the key pluripotency gene’s translation through occupying the particular sites of RNAs. Although the mechanism is unknown, the toxic effects of nCoVN are clear, which reminds us that SARS-CoV-2 might impair the reproductive system and hematopoietic system. In conclusion, we first reported expressing nCoVN could totally change the cell fate of iPSC, which provided new clues to help people fighting against the virus.

## 6 Author Contributions

ZL, ZW, PW, and BL had substantial contributions to the design of the paper; ZL, ZW, JM, LZ, YQ, and TC performed the experiments and analysed the data; ZC provided critical suggestions to improve the paper; ZL, ZW, PW, and BL wrote the manuscript. All authors (ZL, ZW, JM, LZ, YQ, TC, ZC, PW, and BL) had read and approved the final manuscript.

## 8 Acknowledgments

This work is dedicated to all the medical staff who are still fighting against COVID-19 in China. Your efforts make us safer.

