## Supplementary Figures for "The Nucleocapsid Protein of SARS-CoV-2 Abolished Pluripotency in Human Induced Pluripotent Stem Cells"

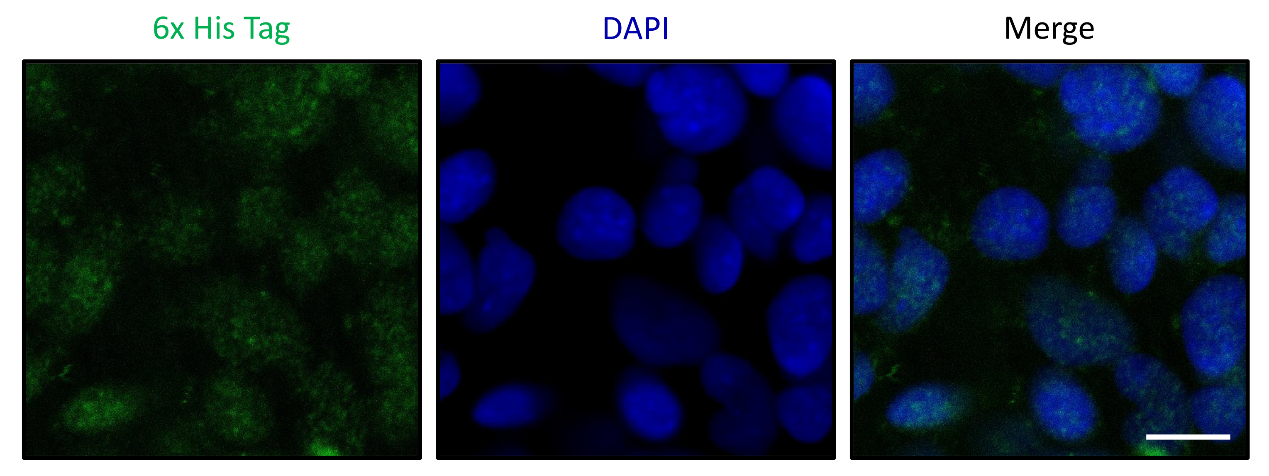

**Supplementary Figure 1**. nCoVN was expressed in the iPSC-nCoVN. Due to a 6× His epitope Tag was fused to the N-terminal of nCoVN, the signal of 6× His Tag could be used to evaluate the expression of nCoVN. Representative immunofluorescent staining images of 6× His Tag (green) in iPSC-nCoVN were shown in the figure. The cell nuclei were stained by DAPI (blue). The scale bar represents 10 μm.

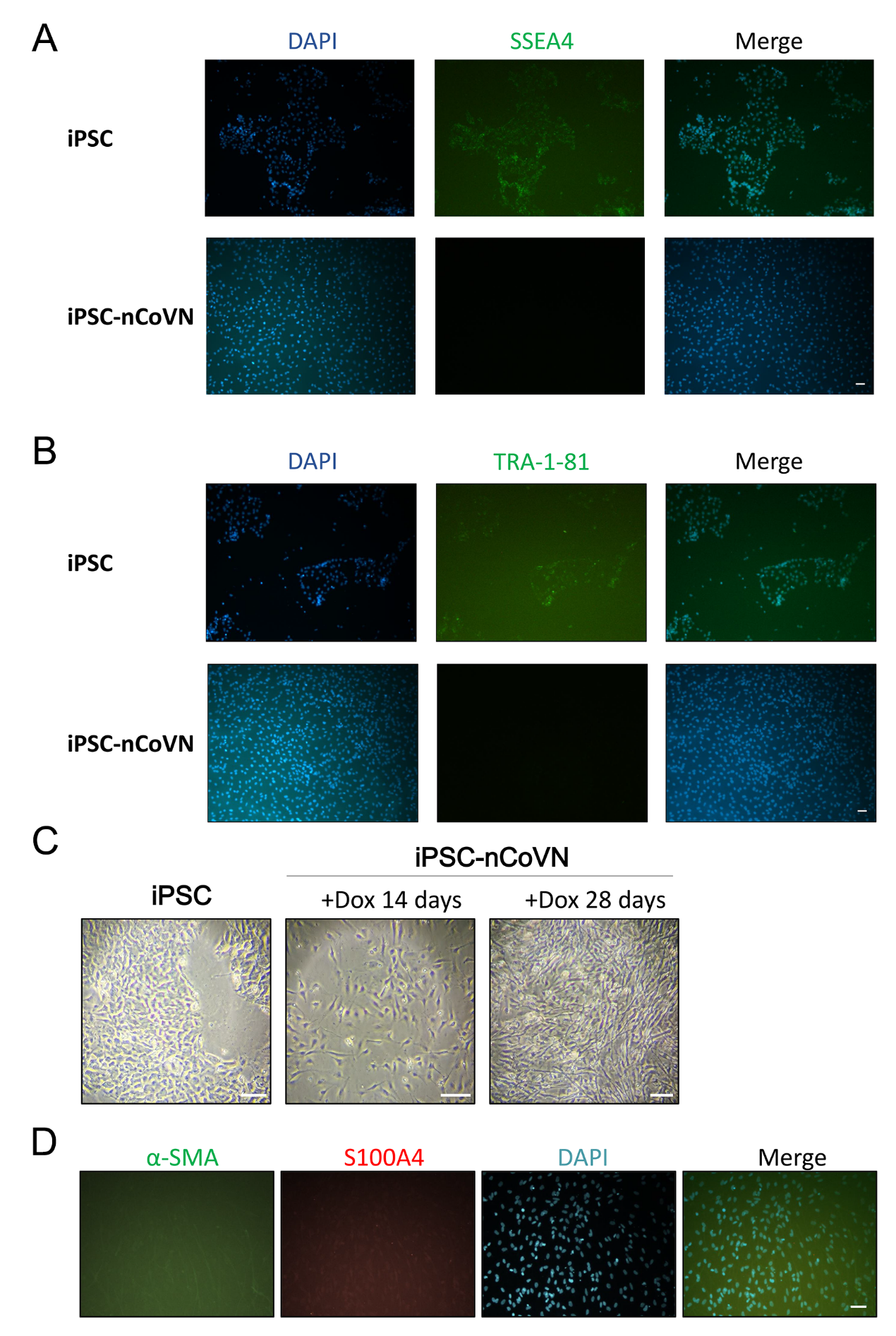

**Supplementary Figure 2**. Expression of nCoVN changed the pluripotent fate of iPSC. (A-B) Representative immunofluorescent staining images of pluripotency markers (A) SSEA4 (green) and, (B) TRA-1-81 (green) in iPSC and iPSC-nCoVN after a 14-day induction. The cell nuclei were stained by DAPI (blue). Scale bars represent 50 μm. (C) The morphology of iPSC, iPSC-nCoVN under a 14-day induction, and iPSC-nCoVN under a 28-day induction. The scale bar is 100 μm. (D) Representative immunofluorescent staining images of fibroblast markers alpha-smooth muscle actin (α-SMA, green) and S100A4 (red) in iPSC-nCoVN after a 28-day induction. The cell nuclei were stained by DAPI (blue). iPSC-nCoVN exhibited fibroblast-like morphology. The scale bar represents 50 μm.
